# Roles of lysosomotropic agents on LRRK2 activation and Rab10 phosphorylation

**DOI:** 10.1101/2020.08.25.267385

**Authors:** Tomoki Kuwahara, Kai Funakawa, Tadayuki Komori, Maria Sakurai, Gen Yoshii, Tomoya Eguchi, Mitsunori Fukuda, Takeshi Iwatsubo

## Abstract

Leucine-rich repeat kinase 2 (LRRK2), the major causative gene product of autosomal-dominant Parkinson’s disease, is a protein kinase that phosphorylates a subset of Rab GTPases. Since pathogenic LRRK2 mutations increase its ability to phosphorylate Rab GTPases, elucidating the mechanisms of how Rab phosphorylation is regulated by LRRK2 is of great importance. We have previously reported that chloroquine-induced lysosomal stress facilitates LRRK2 phosphorylation of Rab10 to maintain lysosomal homeostasis. Here we reveal that Rab10 phosphorylation by LRRK2 is potently stimulated by treatment of cells with a set of lysosome stressors and clinically used lysosomotropic drugs. These agents commonly promoted the formation of LRRK2-coated enlarged lysosomes and extracellular release of lysosomal enzyme cathepsin B, the latter being dependent on LRRK2 kinase activity. In contrast to the increase in Rab10 phosphorylation, treatment with lysosomotropic drugs did not increase the enzymatic activity of LRRK2, as monitored by its autophosphorylation at Ser1292 residue, but rather enhanced the molecular proximity between LRRK2 and its substrate Rab GTPases on the cytosolic surface of lysosomes. Lysosomotropic drug-induced upregulation of Rab10 phosphorylation was likely a downstream event of Rab29 (Rab7L1)-mediated enzymatic activation of LRRK2. These results suggest a regulated process of Rab10 phosphorylation by LRRK2 that is associated with lysosomal overload stress, and provide insights into the novel strategies to halt the aberrant upregulation of LRRK2 kinase activity.

## Introduction

Parkinson’s disease (PD) is a major neurodegenerative disease of the adulthood mainly affecting the extrapyramidal motor system. The Leucine-rich repeat kinase 2 (*LRRK2*) gene has been identified as the most common causative gene for autosomal dominant form of familial PD (Paisan-Ruiz et al., 2004; Zimprich et al., 2004). *LRRK2* gene has also been known as a risk factor of other diseases affecting non-neuronal organs, such as Crohn’s disease or Leprosy (Barrett et al., 2008; Zhang et al., 2009). LRRK2 encodes a large protein kinase that harbors multiple characteristic domains, and its pathogenic mutations associated with PD span multiple domains. The expression pattern of LRRK2 *in vivo* is relatively ubiquitous, but it is noteworthy that macrophages, neutrophils and other immune cells display high expression of LRRK2 (Fan et al., 2018; Gardet et al., 2010; Hakimi et al., 2011; Thevenet et al., 2011), suggesting the diverse roles of LRRK2 such as immune-related functions. Intracellular functions of LRRK2 have been considered to be linked to the regulation of vesicle trafficking around endolysosomes (Cookson, 2016), and previous studies on *Lrrk2* knockout rodents have shown the accumulation of enlarged lysosomes or related organelles in the kidney and lung (Baptista et al., 2013; Fuji et al., 2015; Herzig et al., 2011; Hinkle et al., 2012; Tong et al., 2010). These observations led us to hypothesize that LRRK2 plays a physiologically pivotal role in the maintenance of lysosome morphology and functions.

The role of LRRK2 as a protein kinase has long been unknown, but it has been acknowledged that autophosphorylation at Ser1292 residue serves as an indicator of LRRK2 kinase activity (Sheng et al., 2012). It has also been known that the phosphorylation at residues Ser935 or Ser910 in LRRK2 is rapidly decreased upon treatment with LRRK2 kinase inhibitors (Deng et al., 2011; Dzamko et al., 2010; Reith et al., 2012), although this phosphorylation is not an autophosphorylation (Dzamko et al., 2010) and is not always correlated with the kinase activity of LRRK2 (Christensen et al., 2018; Ito et al., 2014). In 2016, a subset of Rab GTPases were identified as *bona fide* cellular substrates of LRRK2 (Steger et al., 2016). The well-studied substrates are Rab8 and Rab10, and the phosphorylation site is located in a well-conserved switch II region of Rab GTPases, i.e., Thr72 residue of Rab8a and Thr73 residue of Rab10. Remarkably, most of the typical PD-associated LRRK2 mutations elicit a dramatic increase in the phosphorylation of these substrate Rab GTPases. The activation of LRRK2 has also been reported in neurons in sporadic PD brains (Di Maio et al., 2018; Fraser et al., 2016). These findings on LRRK2 raise the possibility that hyperphosphorylation of Rab GTPases acts as a trigger of neurodegeneration, and thus, inhibiting the activation of LRRK2 is expected to be a promising therapeutic strategy for PD. In fact, LRRK2 kinase inhibitors are being tested in clinical trials (https://clinicaltrials.gov/).

However, it remains largely unclear as to when and how LRRK2 is activated. Expression of Rab29 (also known as Rab7L1), a member of Rab GTPase family encoded in a PD risk locus *PARK16* (Satake et al., 2009), and the introduction of a familial PD-causative mutation in VPS35 (D620N) (Vilariño-Güell et al., 2011) have been reported to prompt LRRK2 activation (Liu et al., 2018; Mir et al., 2018; Purlyte et al., 2018). Notably, several studies have demonstrated that Rab29 regulates the subcellular localization as well as the activity of LRRK2; overexpression of Rab29 causes LRRK2 recruitment onto the Golgi apparatus (Madero-Perez et al., 2018; Purlyte et al., 2018), and membrane association is required for the Rab29 activation of LRRK2 (Gomez et al., 2019). We have previously shown that endogenous Rab29 recruits LRRK2 onto the enlarged lysosomal membranes when cells are exposed to chloroquine, a lysosomotropic agent that causes lysosomal stress (Eguchi et al., 2018). This recruitment of LRRK2 then evokes the translocation of its substrates Rab8/10 onto the lysosomal membranes in a phosphorylation-dependent manner, leading to the inhibition of lysosomal enlargement and the exocytic release of lysosomal content. These findings, together with other studies noted above, prompted us to propose a “Rab29-LRRK2-Rab8/10 cascade” in the pathobiology of LRRK2 (Kuwahara and Iwatsubo, 2020). Notably, our results revealed that chloroquine treatment also increases Rab10 phosphorylation at Thr73 by LRRK2. However, it remained unclear whether this increase in Rab phosphorylation is induced by lysosomal overload stresses in a general fashion, or alternatively, the effect is specific to chloroquine. In addition, it remains to be solved whether the activation of Rab phosphorylation by LRRK2 is attributed to the increase in LRRK2 enzymatic activity, or to other mechanisms such as the increased proximity of LRRK2 and Rab caused by the alteration of their subcellular localizations.

Here, we examined the effect of LRRK2 on Rab10 phosphorylation under various stress conditions, employing pharmacological approaches. Our results showed that the induction of Rab10 phosphorylation and upregulation in lysosomal release that are dependent on LRRK2 kinase activity were common effects of lysosomotropic drugs. Notably, the induction of Rab10 phosphorylation was attributable to the enhanced proximity of LRRK2 and Rab10, which occurred downstream of the Rab29-mediated enzymatic activation of LRRK2. Taken together, these results offer a clearer insight into the regulatory mechanisms of Rab10 phosphorylation by LRRK2.

## Results

### Pharmacological induction of lysosomal stress, but not other types of stresses, activates Rab10 phosphorylation

We have previously reported that LRRK2-mediated phosphorylation of Rab10 at Thr73 residue increased upon treatment of cells with chloroquine, a representative lysosomal stressor having lysosomotropic property (Eguchi et al., 2018). Thus, we first examined whether other cellular stresses induced by pharmacological treatments also upregulate Rab10 phosphorylation. To this end, we selected a set of compounds that have been widely used to induce cellular stresses in a variety of experimental paradigms. The lysosomal stressors include bafilomycin A1 (vacuolar proton pump inhibitor), ammonium chloride (NH4Cl; lysosomotropic weak base) and monensin (Na^+^/H^+^ ionophore that results in an increase in lysosomal osmotic pressure) (Florey et al., 2015) in addition to chloroquine, all of which inhibit lysosomal acidification (Florey et al., 2015; Misinzo et al., 2008). We also included compounds that elicit other types of cellular stresses, i.e., intracellular vacuole formation (vacuolin-1), macroautophagy (Torin 1), mitophagy (carbonyl cyanide m-chlorophenylhydrazone; CCCP), ER stress (Tunicamycin), oxidative stress (Rotenone) and inflammation (lipopolysaccharide; LPS). RAW264.7 cells, a mouse macrophage cell line, were first pretreated for 48 hrs with interferon-γ (IFN-γ), a well-known procedure to drive endogenous LRRK2 expression (Gardet et al., 2010) and then treated with the reagents above for 3 hrs at concentrations that were considered sufficient to exert pharmacological effects. Rab10 phosphorylation in cells were then examined by immunoblot analysis with an anti-phopho-Thr73 Rab10 antibody. We found that Rab10 phosphorylation was potently induced by chloroquine or monensin (*p* < 0.0001), and moderately by bafilomycin A1 or NH4Cl (*p* < 0.01) **(Fig. 1A and 1B)**. On the other hand, treatment with other compounds had little or no effect on the phosphorylation state of Rab10 **(Fig. 1A and 1B)**. Similar changes in the phosphorylation state were observed using an anti-phospho-Thr72 Rab8 antibody, although this antibody crossreacts with phosphorylated Rab3A, Rab10, Rab35 and Rab43 (source: information provided by the manufacturer on the datasheet) **(Fig. 1A)**. Treatment with a LRRK2-specific kinase inhibitor GSK2578215A (Reith et al., 2012) inhibited the drug-induced increase in Rab10 phosphorylation **(Fig. 1A and 1B)**. The effectiveness of LRKR2 inhibitor was further confirmed by the reduction in the phospho-Ser935 signal in LRRK2 that has been established as a marker of LRRK2 kinase inhibition (Deng et al., 2011; Dzamko et al., 2010; Reith et al., 2012) **(Fig. 1A)**. The phospho-Ser935 signal was exceptionally increased by LPS treatment, which was likely due to stimulation of the Toll-like receptor signaling (Dzamko et al., 2012). Protein levels of LRRK2 and α-tubulin were not affected upon exposure to these stressors **(Fig. 1A)**. Because chloroquine and monensin both have a strong potency to increase the lysosomal osmotic pressure, we considered that lysosomal stresses, especially those causing an overload state, selectively induced the upregulation of Rab10 phosphorylation in cells.

**Fig. 1.**
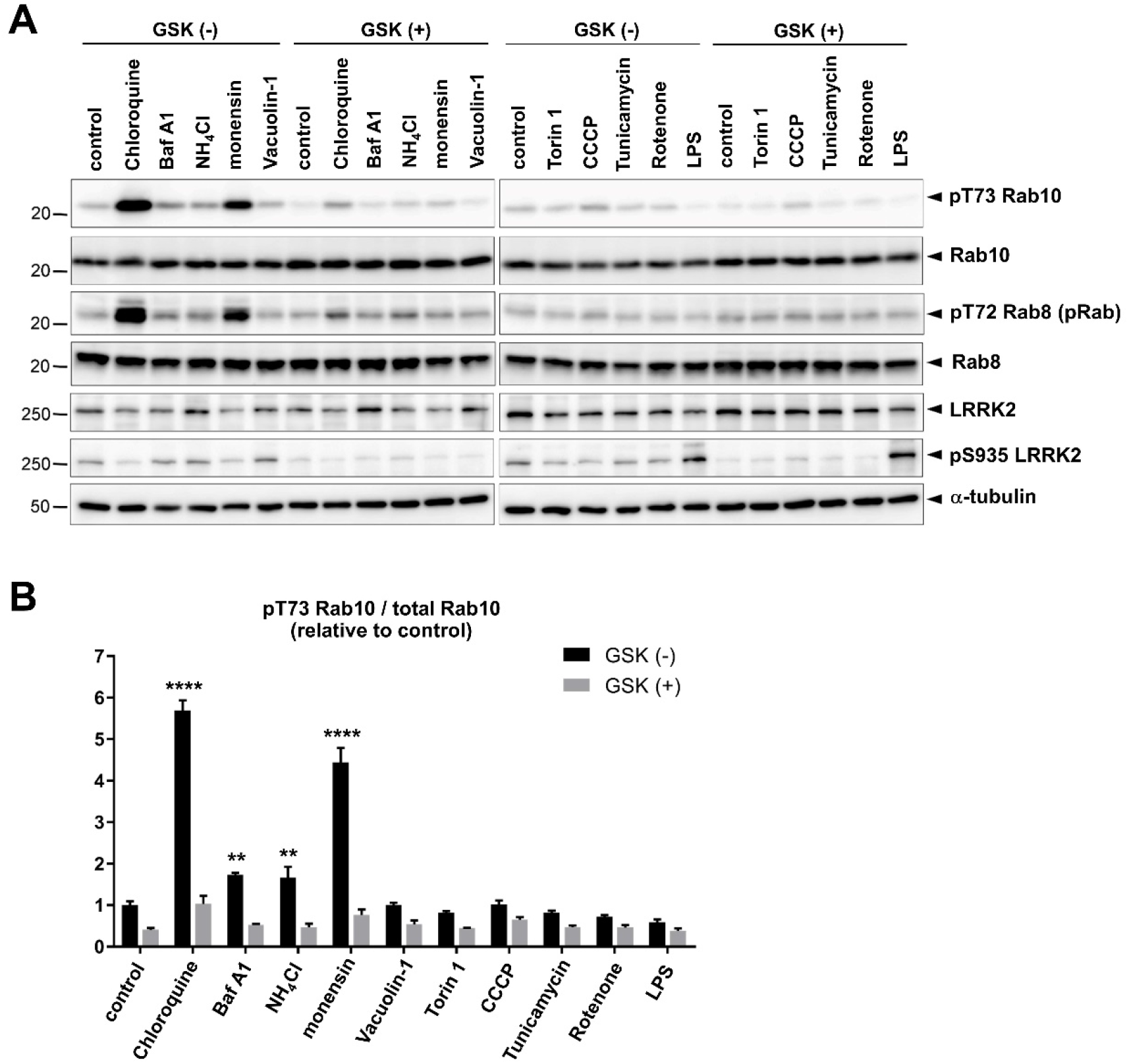
Pharmacological induction of lysosomal stress, but not other types of stresses, activates Rab10 phosphorylation. **(A)** Immunoblot analysis of lysates from RAW264.7 cells treated for 3 hrs with the indicated reagents with or without LRRK2 kinase inhibitor GSK2578215A (GSK), as indicated. Representative pictures analyzed for phospho-Thr73-Rab10, Rab10, phospho-Thr72-Rab8, Rab8, LRRK2, phospho-Ser935-LRRK2 and α-tubulin are shown. **(B)** Densitometric analysis of Rab10 phosphorylation, as shown in A. The levels of phospho-Rab10 were normalized by those of Rab10. Relative values compared to control-GSK (-) were shown. Data represent mean ± SEM. n = 4 for GSK (-), n = 3 for GSK (+), \*\**P* < 0.01, \*\*\*\**P* < 0.0001 compared with control GSK (-), two-way ANOVA with Dunnett’s test.

### Clinically used lysosomotropic drugs activate LRRK2 phosphorylation of Rab10

Next, we sought to examine whether the stimulation of LRRK2 phosphorylation of Rab10 is a common feature of small lysosomotropic compounds causing lysosomal overload. Generally, weak base compounds having lipophilic or amphiphilic property are readily protonated and trapped within lysosomes, causing lysosomal overload and swelling (de Duve et al., 1974; Kuzu et al., 2017; Marceau et al., 2012; Villamil Giraldo et al., 2014) **(Fig. 2A)**. There exist a set of clinically used compounds that are known to exhibit lysosomotropism, including antipsychotic and anticancer drugs. We thus selected 11 of the commonly used clinical drugs having such lysosomotropic properties, as listed in **Table 1**. Among these, eight drugs (chloroquine, chlorpromazine, thioridazine, nortriptyline, fluoxetine, imanitib, gefitinib and tamoxifen) possess a relatively high basicity (pKa, the logarithm of the dissociation constant of the most basic center of the compound) and lipophilicity (ClogP, the partition coefficient of the neutral species of a compound between octanol and water), whereas other three drugs (lidocaine, procainamide and nifedipine) possess hydrophilic property and thus require higher doses for entry into lysosomes **(Fig. 2B)**; these drugs are classified as Class-II and Class-I lysosomotropic compounds, respectively (Kuzu et al., 2017). RAW264.7 cells were exposed to these drugs for 3 hrs at concentrations equal to or slightly exceeding the range of those causing a decrease in LysoTracker fluorescence **(Table 1)**. Immunoblot analysis revealed that the phosphorylation of Rab10 was induced by treatment with 7 different drugs: chloroquine, chlorpromazine, thioridazine, nortriptyline, gefitinib, tamoxifen and lidocaine **(Fig. 2C and 2D)**. Treatment with other drugs – fluoxetine, imanitib, procainamide and nifedipine – also showed a trend of increased Rab10 phosphorylation which, however, did not reach a statistical significance. Additional treatment with LRRK2 kinase inhibitor GSK2578215A almost completely inhibited the drug-induced increase in Rab10 phosphorylation **(Fig. 2C and 2D)**. The effectiveness of the LRKR2 kinase inhibitor was again confirmed by the reduction in phospho-Ser935 signal of LRRK2 **(Fig. 2C)**. These data support the notion that a set of clinically used lysosomotropic drugs have the potential to stimulate LRRK2-mediated phosphorylation of Rab10.

**Fig. 2.**
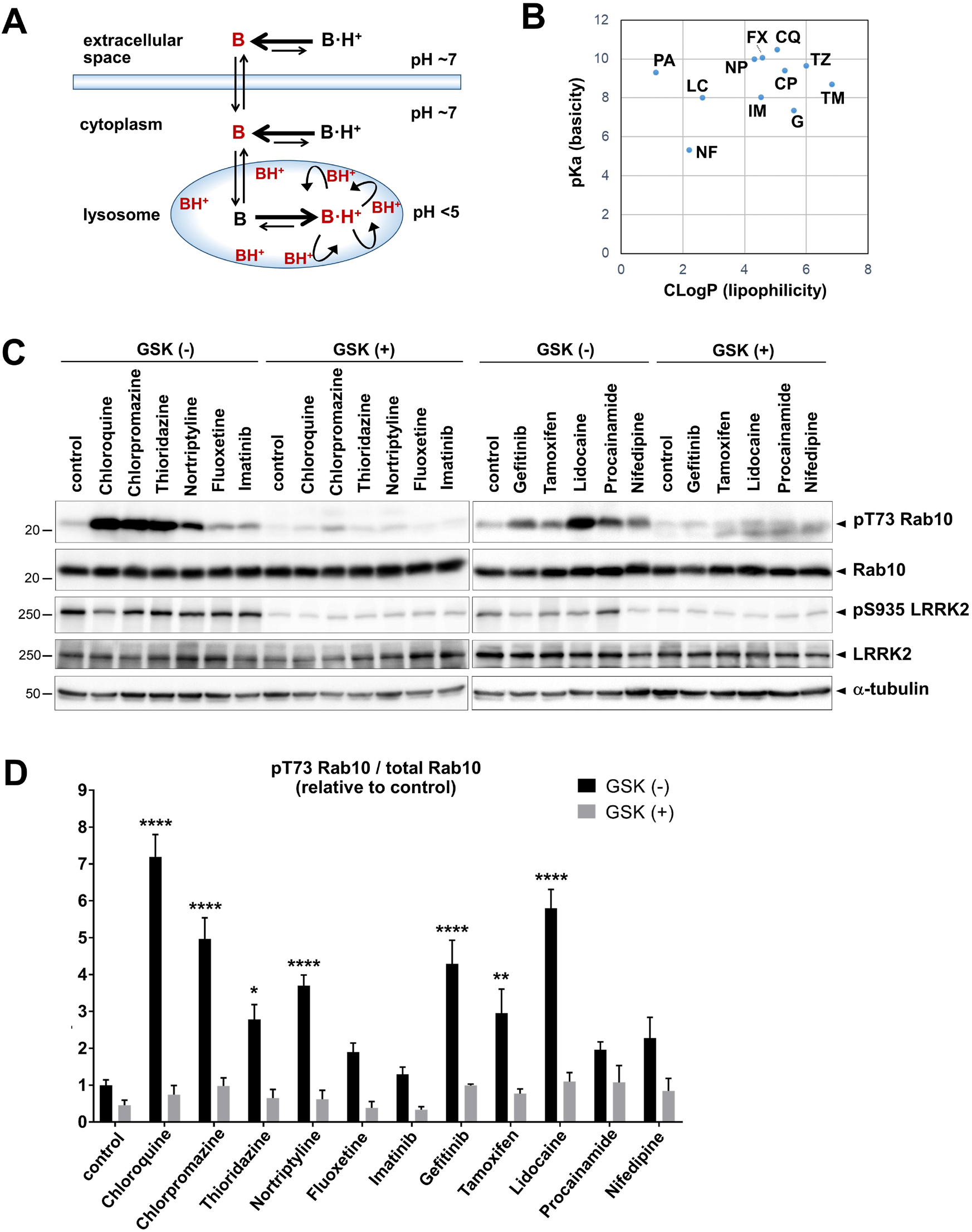
Clinically used lysosomotropic drugs activate LRRK2 phosphorylation of Rab10. **(A)** The cation trapping mechanism of lysosomotropic agents inside lysosomes. Weak base compounds passively diffuse through cell membranes and reach acidic environment of lysosomes, where they get charged and can no longer pass through the lysosomal membranes. **(B)** A scatter plot showing the distribution of lysosomotropic drugs used in this study. The basicity (pKa value) and lipophilicity (ClogP value) of each drug are shown. For abbreviations of drug names, see Table 1. **(C)** Immunoblot analysis of lysates from RAW264.7 cells treated for 3 hrs with a set of clinically used lysosomotropic drugs with or without LRRK2 kinase inhibitor GSK2578215A (GSK), as indicated. Representative pictures analyzed for phospho-Thr73-Rab10, Rab10, phospho-Ser935-LRRK2, LRRK2 and α-tubulin are shown. **(D)** Densitometric analysis of Rab10 phosphorylation, as shown in C. The levels of phospho-Rab10 were normalized by those of Rab10. Relative values compared to control-GSK (-) were shown. Data represent mean ± SEM. n = 10 for GSK (-), n =4 for GSK (+), \**P* < 0.05, \*\**P* < 0.01, \*\*\*\**P* < 0.0001 compared with control GSK (-), two-way ANOVA with Dunnett’s test.

**Table 1.**
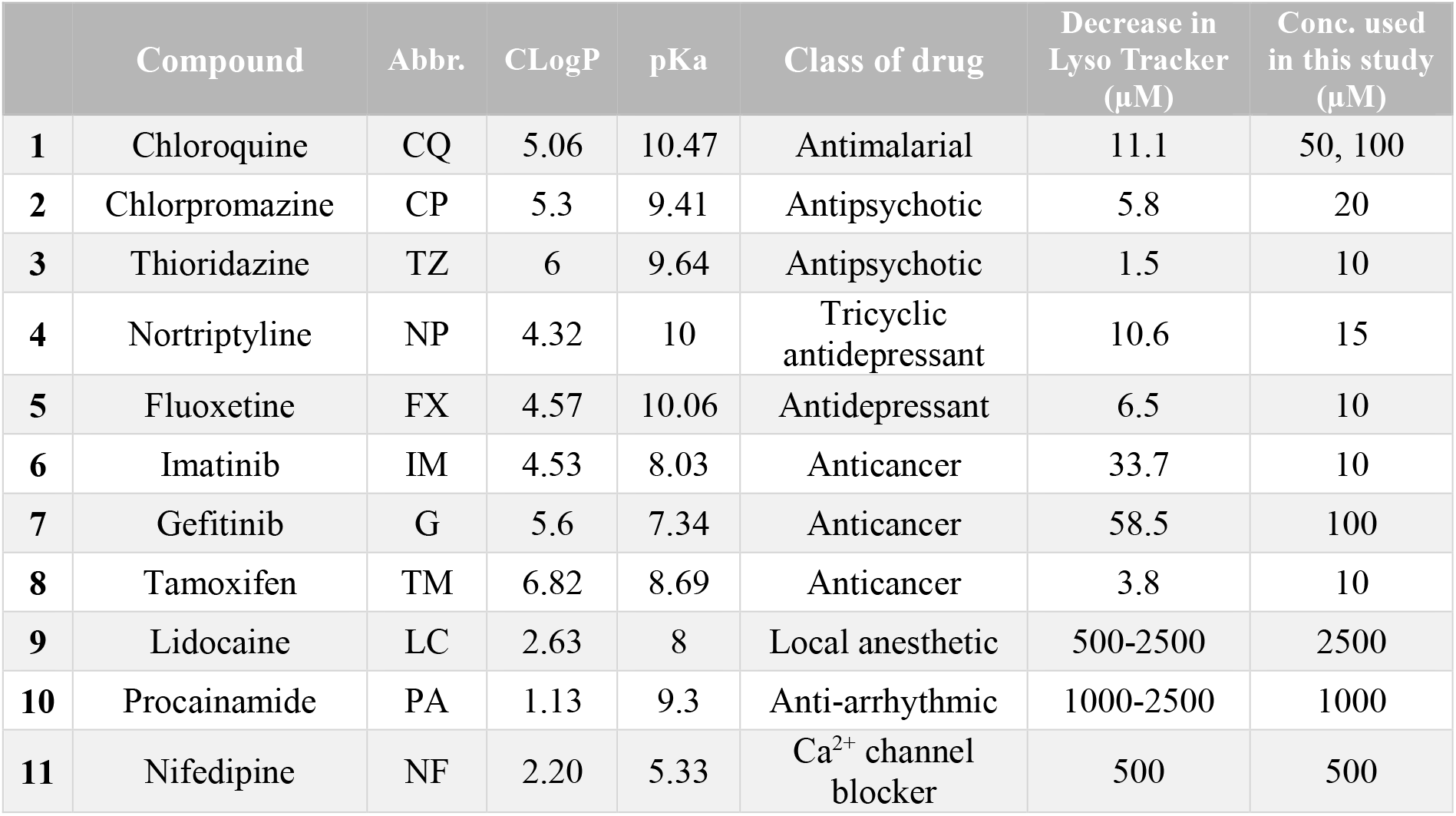
Clinically used lysosomotropic drugs tested in this study.

|  | Compound | Abbr. | CLogP | pKa | Class of drug | Decrease in Lyso Tracker (μM) | Conc. used in this study (μM) |
| --- | --- | --- | --- | --- | --- | --- | --- |
| 1 | Chloroquine | CQ | 5.06 | 10.47 | Antimalarial | 11.1 | 50, 100 |
| 2 | Chlorpromazine | CP | 5.3 | 9.41 | Antipsychotic | 5.8 | 20 |
| 3 | Thioridazine | TZ | 6 | 9.64 | Antipsychotic | 1.5 | 10 |
| 4 | Nortriptyline | NP | 4.32 | 10 | Tricyclic antidepressant | 10.6 | 15 |
| 5 | Fluoxetine | FX | 4.57 | 10.06 | Antidepressant | 6.5 | 10 |
| 6 | Imatinib | IM | 4.53 | 8.03 | Anticancer | 33.7 | 10 |
| 7 | Gefitinib | G | 5.6 | 7.34 | Anticancer | 58.5 | 100 |
| 8 | Tamoxifen | TM | 6.82 | 8.69 | Anticancer | 3.8 | 10 |
| 9 | Lidocaine | LC | 2.63 | 8 | Local anesthetic | 500-2500 | 2500 |
| 10 | Procainamide | PA | 1.13 | 9.3 | Anti-arrhythmic | 1000-2500 | 1000 |
| 11 | Nifedipine | NF | 2.20 | 5.33 | Ca<sup>2+</sup> channel blocker | 500 | 500 |

### Lysosomotropic drugs induce the formation of LRRK2-positive enlarged lysosomes and LRRK2-dependent exocytic release of cathepsin B

It is well known that lysosomotropic agents induce massive cytoplasmic vacuolation that have characteristics of lysosomal properties (Kuzu et al., 2017; Marceau et al., 2012). We have also previously shown that the enlarged lysosomes induced by chloroquine are coated with LRRK2 (Eguchi et al., 2018). Thus, we examined whether other lysosomotropic drugs can induce the enlargement of lysosomes as well as the LRRK2 recruitment. Confocal microscopic analyses indicated that the treatment either with chloroquine, chlorpromazine, nortriptyline or gefitinib similarly induced the accumulation of endogenous LRRK2 in LAMP1-positive enlarged lysosomes **(Fig. 3A and 3B)**, in accordance with our prediction.

**Fig. 3.**
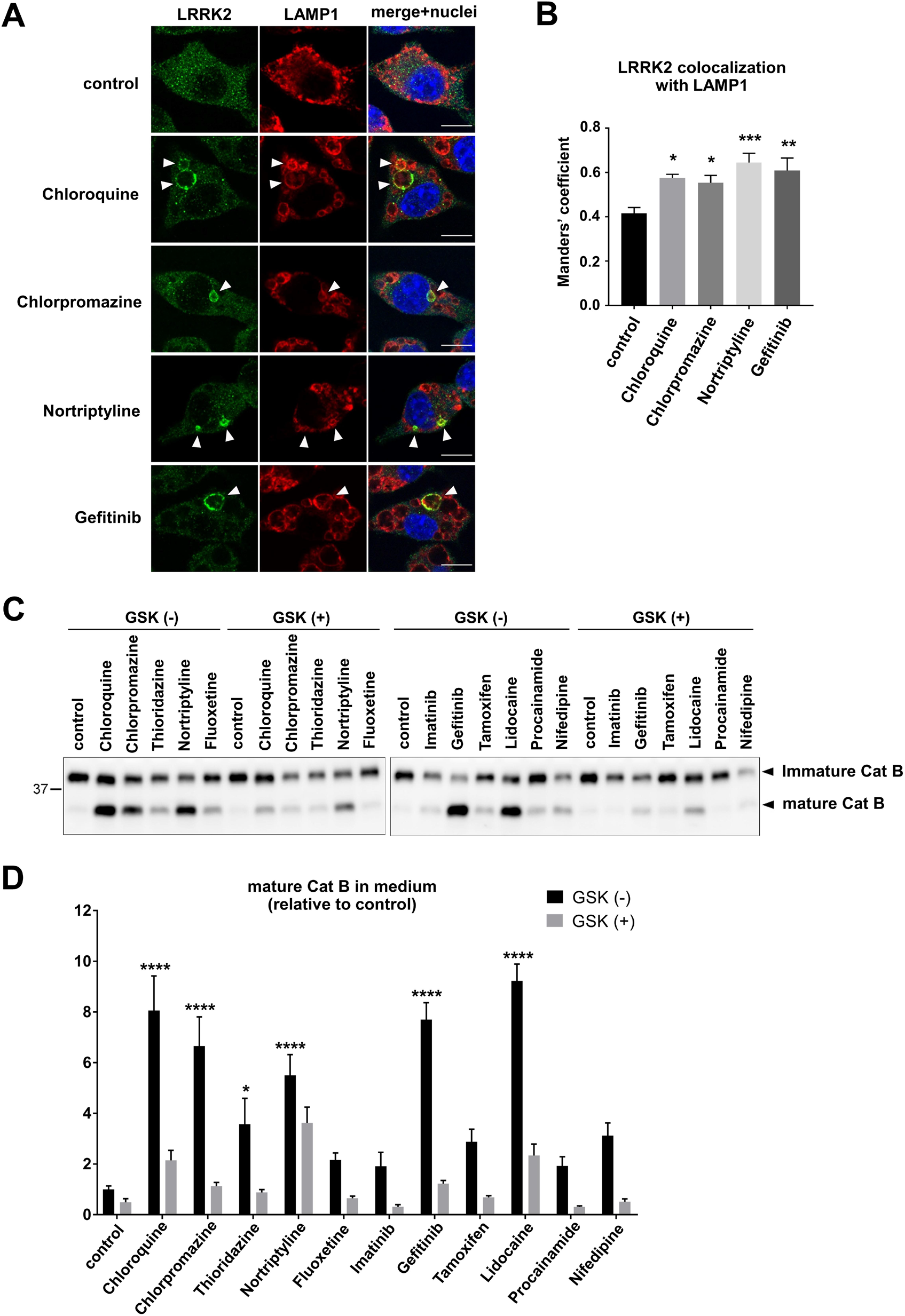
Lysosomotropic drugs induce the formation of LRRK2-positive enlarged lysosomes and release of cathepsin B. **(A)** Immunocytochemical analysis of lysosomal localization of endogenous LRRK2 (pointed by arrowheads) in RAW264.7 cells treated with the indicated drugs. Scale bars = 10 μm. **(B)** Quantitative measurements of colocalization of LRRK2 with LAMP1, as shown in A, using Manders’ coefficient. Data represent mean ± SEM. n = 15, \**P* < 0.05, \*\**P* < 0.01, \*\*\**P* < 0.001 compared with control, two-way ANOVA with Dunnett’s test. **(C)** Immunoblot analysis of the levels of immature and mature cathepsin B (Cat B) in media of RAW264.7 cells treated for 3 hrs with the indicated drugs with or without LRRK2 kinase inhibitor GSK2578215A (GSK). Representative pictures analyzed for Cat B are shown. **(D)** Densitometric analysis of Cat B release, as shown in C. The levels of mature Cat B in media were normalized by those of α-tubulin in cell lysates. Relative values compared to control-GSK (-) were shown. Data represent mean ± SEM. n = 6 for GSK (-), n =4 for GSK (+), \**P* < 0.05, \*\*\*\**P* < 0.0001 compared with control GSK (-), two-way ANOVA with Dunnett’s test.

We have previously shown that chloroquine treatment in RAW264.7 cells caused rapid exocytic release of lysosomal content, i.e., cathepsin B and cathepsin D, through LRRK2-mediated phosphorylation of Rab10 (Eguchi et al., 2018). We thus tested whether other lysosomotropic drugs similarly induce the release of cathepsin B by LRRK2. RAW264.7 cells were exposed to 11 different drugs **(Table 1)** for 3 hrs as used in Fig. 1, then culture media of drug-treated cells were collected. Immunoblot analysis indicated that the release of cathepsin B was significantly promoted by treatment with chloroquine, chlorpromazine, thioridazine, nortriptyline, gefitinib and lidocaine **(Fig. 3C and 3D)**. This set of drugs almost coincided with those that induced Rab10 phosphorylation. The relative potency of these drugs to induce cathepsin B release was proportional to that on Rab10 phosphorylation, as chloroquine and chlorpromazine had potent effects on inducing both Rab10 phosphorylation and cathepsin B release, whereas thioridazine had moderate effects. The release of cathepsin B was not likely due to cell death, as supported by the unaltered levels of lactate dehydrogenase (LDH) released into media **(Supplementary Fig. 1)** and of α-tubulin in cell lysates **(Fig. 2C)**. To confirm the dependency on LRRK2 kinase activity, cells were treated with a LRRK2 inhibitor GSK2578215A in addition to lysosomotropic drugs similarly to the experiments in Fig. 2C-2D. The release of cathepsin B was significantly inhibited by treatment with GSK2578215A **(Fig. 3C and 3D)**, although the inhibitory effect was not necessarily complete. These results collectively suggest that lysosomotropism contributes to a common cellular mechanism that stimulates LRRK2 phosphorylation of Rab10, LRRK2 recruitment onto lysosomes and LRRK2-mediated release of lysosomal content.

### Chloroquine induces Rab10 phosphorylation in various types of cells without increasing the enzymatic activity of LRRK2

We next examined whether the induction of LRRK2 phosphorylation of Rab10 is commonly observed across various cell types and primary cells in addition to IFN-γ-treated RAW264.7 cells that express high levels of LRRK2. We then focused our attention on chloroquine as a lysosomotropic drug that showed the highest effects on Rab10 phosphorylation and cathepsin release. Treatment of either mouse 3T3-Swiss albino fibroblasts or mouse bone marrow-derived primary macrophages with chloroquine caused potent induction of Rab10 phosphorylation **(Fig. 4A-4C)**. Chloroquine treatment of human embryonic kidney (HEK) 293 cells and human neuroblastoma SH-SY5Y cells, both of which express relatively low levels of endogenous LRRK2, caused a modest increase in Rab10 phosphorylation, and exogenous expression of pathogenic mutant LRRK2 (G2019S) further increased Rab10 phosphorylation **(Fig. 4D-4I)**.

**Fig. 4.**
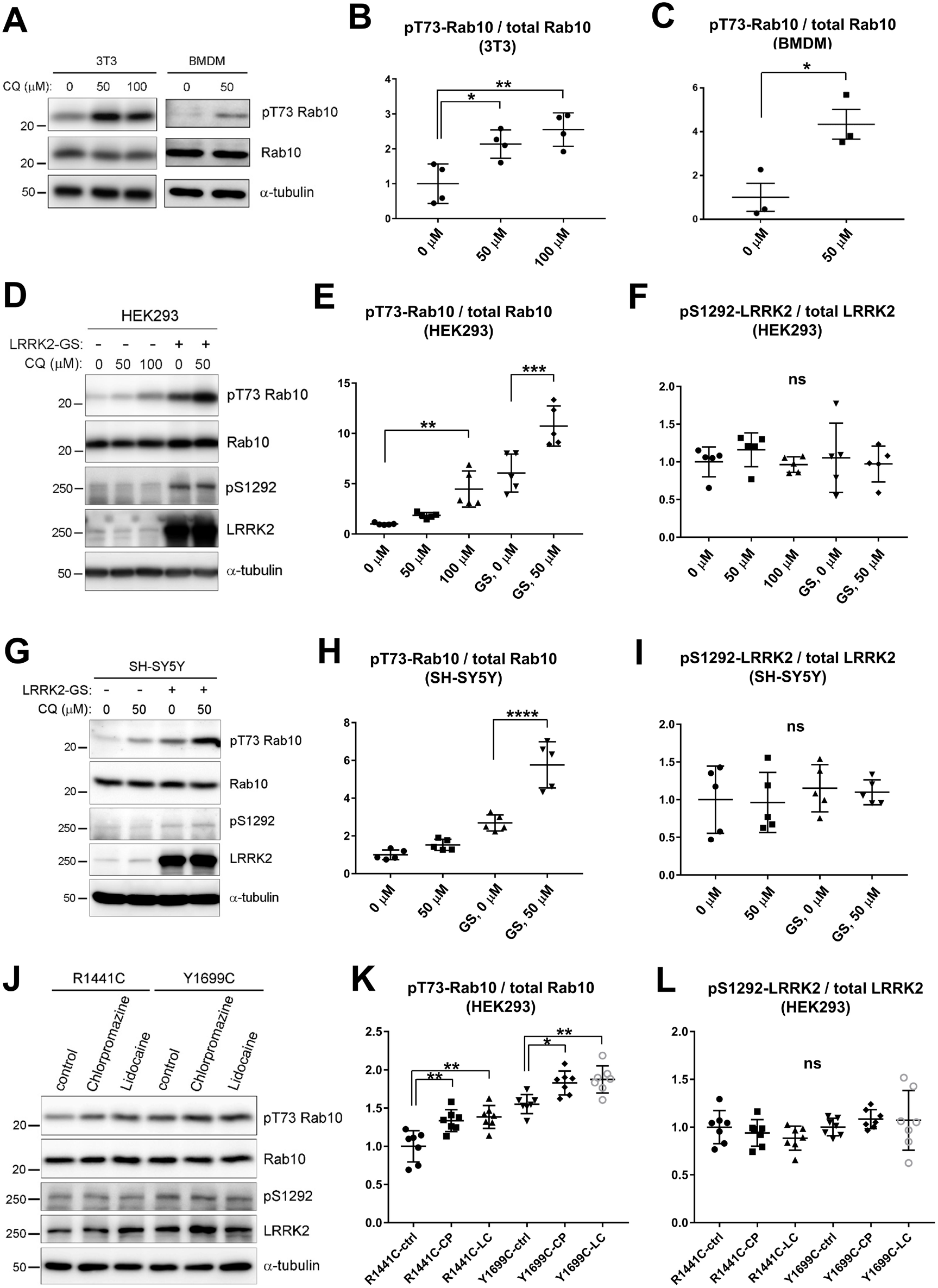
Chloroquine treatment induces Rab10 phosphorylation without increasing the enzymatic activity of LRRK2. **(A)** Immunoblot analysis of the increased Rab10 phosphorylation upon treatment with chloroquine in 3T3-Swiss albino cells and mouse bone marrow-derived primary macrophages (BMDM). Representative pictures are shown. **(B, C)** Densitometric analysis of the level of phospho-pThr73-Rab10 normalized by that of Rab10 in 3T3 cells (B) and BMDM (C) exposed to chloroquine, as shown in A. Relative values compared to control (0 μM) were shown. Data represent mean ± SD. n = 4 (B) or n = 3 (C), \**P* < 0.05, \*\**P* < 0.01 compared with control (0 μM), by one-way ANOVA with Dunnett’s test (B) or by Student *t* test (C). **(D)** Immunoblot analysis of the phosphorylations of Rab10-Thr73 and LRRK2-Ser1292 upon treatment with chloroquine in HEK293 cells with or without overexpression of G2019S mutant LRRK2 (LRRK2-GS). Representative pictures are shown. **(E, F)** Densitometric analysis of the levels of phospho-Thr73-Rab10 normalized by that of Rab10 (E) or phospho-Ser1292-LRRK2 normalized by that of LRRK2 (F) in HEK293 cells exposed to chloroquine, as shown in D. Relative values compared to control (0 μM) were shown. Data represent mean ± SD. n = 5 each, \*\**P* < 0.01, \*\*\**P* < 0.001, ns: not significant, one-way ANOVA with Dunnett’s test. **(G)** Immunoblot analysis of the phosphorylations of Rab10-Thr73 and LRRK2-Ser1292 upon treatment with chloroquine in SH-SY5Y cells with or without overexpression of LRRK2-GS. **(H, I)** Densitometric analysis of the level of phospho-Thr73-Rab10 normalized by that of Rab10 (H) or the level of phospho-Ser1292-LRRK2 normalized by that of LRRK2 (I) in SH-SY5Y cells exposed to chloroquine, as shown in G. Relative values compared to control (0 μM) were shown. Mean ± SD, n = 5 each, \*\*\*\**P* < 0.0001, ns: not significant, one-way ANOVA with Dunnett’s test. **(J)** Immunoblot analysis of the phosphorylation of Rab10-Thr73 and LRRK2-Ser1292 upon treatment with chlorpromazine or lidocaine in HEK293 cells overexpressing either R1441C or Y1699C mutant LRRK2. **(K, L)** Densitometric analysis of the levels of phospho-Thr73-Rab10 normalized by that of Rab10 (K) or phospho-Ser1292-LRRK2 normalized by that of LRRK2 (L) in mutant LRRK2-expressing HEK293 cells exposed to chlorpromazine (CP) or lidocaine (LC), as shown in J. Relative values compared to R1441C-control were shown. Mean ± SD, n = 7 each, \**P* < 0.05, \*\**P* < 0.01, ns: not significant, one-way ANOVA with Tukey’s test.

The activation state of LRRK2 kinase in cells can be monitored by assessing its autophosphorylation at Ser1292 residue (Sheng et al., 2012). We therefore examined whether the addition of chloroquine induces the enzymatic activation of LRRK2 by utilizing an anti-phospho-Ser1292 antibody. Because this antibody exclusively recognizes the phosphorylation of human LRRK2, we examined HEK293 and SH-SY5Y cells of human origin. Unexpectedly, in contrast to the increase in Rab10 phosphorylation, the phosphorylation at Ser1292 of LRRK2 was not altered in cells after exposure to chloroquine, even under an overexpression condition of human LRRK2 (G2019S) **(Fig. 4D-4I)**. We additionally tested the effects of other lysosomotropic drugs (chlorpromazine and lidocaine) as well as other pathogenic mutant LRRK2 (R1441C and Y1699C) on the phosphorylation. As expected, phosphorylation of Rab10-Thr73, but not that of LRRK2-Ser1292, was significantly increased upon exposure to either of these drugs in HEK293 cells overexpressing pathogenic mutant LRRK2 **(Fig. 4J-4L)**. These results suggest that lysosomotropic drug-induced increase in Rab10 phosphorylation is mediated by a mechanism other than the upregulation of LRRK2 kinase activity.

### Lysosomotropic drug treatment increases molecular proximity between LRRK2 and its substrate Rab GTPases on lysosomes

To explain the effects of lysosomotropic drugs to increase the Rab10 phosphorylation without alteration in the LRRK2 kinase activity, we hypothesized an alternative mechanism that increases the proximity of LRRK2 and Rab10 in response to lysosomal overload. It has been reported that the membranes of giant vacuoles induced by lysosomotropic agents contain not only lysosomal markers but late endosomal and sometimes *trans*-Golgi markers, suggesting the formation of “hybrid organelles” as a result of increased fusion or transport of organelle membranes (Goldman and Krise, 2010; Kuzu et al., 2017; Marceau et al., 2012; Morissette et al., 2004). We examined whether the formation of such hybrid organelles may result in the lysosomal translocation of substrate Rab GTPases, i.e., Rab10 and Rab8, that normally reside on endosomal membranes. We first showed that the LAMP1-positive enlarged lysosomes induced by chloroquine or gefitinib were partially positive for a late endosome marker, lysobisphosphatidic acid (LBPA) (Kobayashi et al., 1999) **(Fig. 5A)**. However, they were negative for an early endosome marker EEA1 **(Fig. 5A)**, supporting the idea that late endosomal components were specifically incorporated in lysosomal membranes upon their enlargement.

**Fig. 5.**
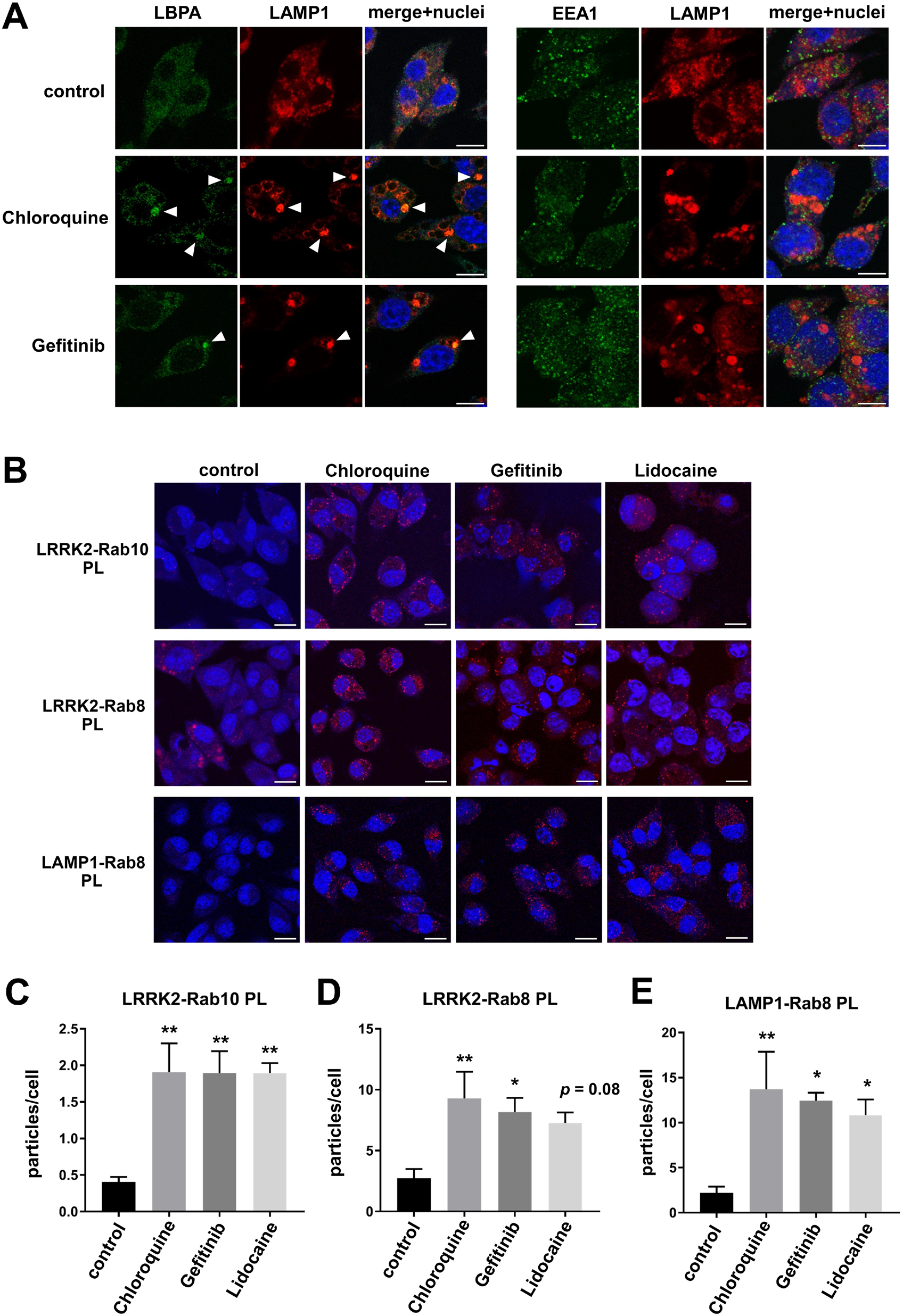
Lysosomotropic drugs increase the molecular proximity between LRRK2 and substrate Rab GTPases on lysosomes. **(A)** Colocalization analysis of LAMP1 with LBPA or EEA1 in RAW264.7 cells exposed to chloroquine, gefitinib or control (PBS). Arrowheads indicate colocalization of both signals. Blue: DRAQ5 (nuclei), scale bars = 10 μm. **(B)** Confocal microscopic detection of proximity ligation (PL) signals (red dots) between LRRK2 and Rab10 (upper panels), LRRK2 and Rab8 (middle panels) or LAMP1 and Rab8 (lower panels) in RAW264.7 cells exposed to the indicated drugs. Blue: DRAQ5 (nuclei), scale bars = 10 μm. **(C-E)** Counting of the particles of PL signals between LRRK2 and Rab10 (C), LRRK2 and Rab8 (D) or LAMP1 and Rab8 (E), as shown in B, using ImageJ software. The total number of particles in the field were divided by that of nuclei to calculate particle number per cell. Data represent mean ± SEM. n = 5, \**P* < 0.05, \*\**P* < 0.01 compared with control, one-way ANOVA with Dunnett’s test.

We then examined whether LRRK2-Rab interaction was increased upon lysosomal enlargement. Because the co-immunoprecipitation analysis failed to detect a stable interaction between LRRK2 and Rab10 even after chloroquine treatment **(Supplementary Fig. 2)**, we analyzed the spatial proximity between LRRK2 and Rab10 or Rab8 upon exposure to lysosomotropic agents by performing *in situ* proximity ligation assay (PLA) (Soderberg et al., 2006; Weibrecht et al., 2010). PLA is a powerful and convenient method that enables direct, sensitive and specific detection of endogenous protein interactions at single-molecule resolution; a key principle is the use of a pair of oligonucleotide-labeled two secondary antibodies (PLA probes) that generate an amplified signal only when the probes are in close proximity (<40 nm). Treatment of RAW264.7 cells with either chloroquine, gefitinib or lidocaine led to a significant increase in proximity ligation (PL) signals between LRRK2-Rab10 as well as LRRK2-Rab8 **(Fig. 5B-5D)**. To further detect the close proximity of these Rabs and lysosomes, we utilized an antibody that recognizes the cytosolic side of LAMP1 that spans lysosomal membranes. We found that, although LAMP1-Rab10 PL signal was absent, LAMP1-Rab8 PL signal was detected and clearly increased upon treatment with either chloroquine, gefitinib or lidocaine **(Fig. 5B and 5E)**. These results indicate the role of lysosomotropic drugs on increasing the molecular proximity between LRRK2 and its substrate Rab GTPases on lysosomes.

### Rab10 phosphorylation by lysosomotropic drugs occurs downstream of Rab29-mediated activation of LRRK2

It has been reported that LRRK2 phosphorylation of Rab10 as well as the LRRK2 enzymatic activity in cells was enhanced by co-expression of Rab29 (also known as Rab7L1) (Liu et al., 2018; Purlyte et al., 2018), another Rab GTPase and a putative PD risk factor encoded in *PARK16* locus (Satake et al., 2009). Rab29 is also known as a genetic and physical interactor of LRRK2 (Beilina et al., 2014; MacLeod et al., 2013) and an upstream regulator that determines the subcellular localization of LRRK2 (Eguchi et al., 2018; Gomez et al., 2019; Liu et al., 2018; Madero-Perez et al., 2018; Purlyte et al., 2018). We therefore examined the relationship between Rab29 and the lysosomotropic agents in the upregulation of Rab10 phosphorylation. When RAW264.7 cells were pretreated with siRNAs against either Rab29 or LRRK2 to suppress their endogenous expression, the induction of Rab10 phosphorylation by chloroquine was significantly inhibited **(Fig. 6A and 6B)**. This suggests that Rab29 is required for the phosphorylation of Rab10 induced by lysosomotropic agents.

**Fig. 6.**
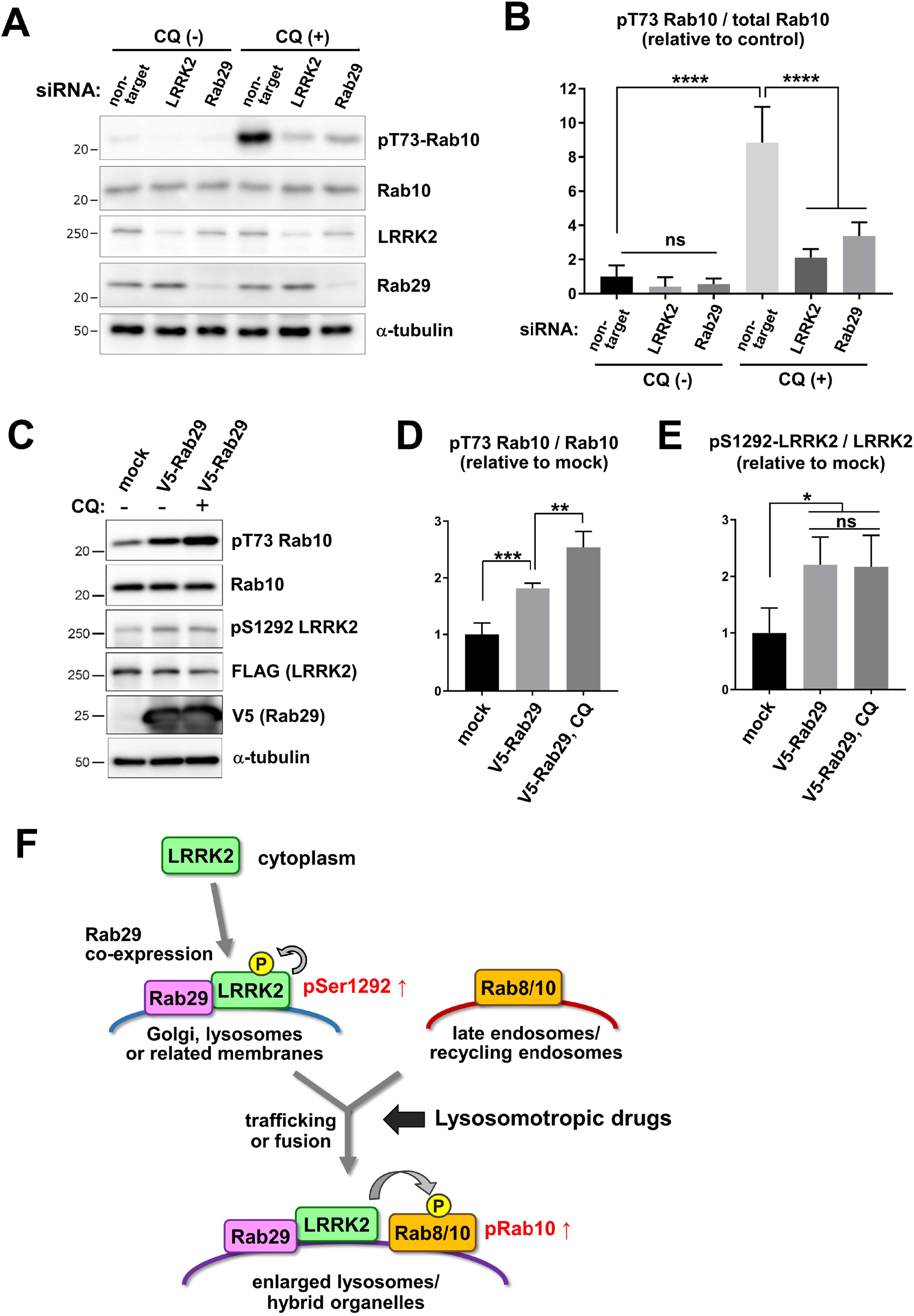
Chloroquine-induced Rab10 phosphorylation occurs downstream of Rab29-mediated activation of LRRK2. **(A)** Immunoblot analysis of lysates from RAW264.7 cells pretreated with siRNA against non-target, LRRK2 or Rab29 and exposed to chloroquine for 3 hrs. Representative pictures analyzed for phospho-Thr73-Rab10, Rab10, LRRK2, Rab29 and α-tubulin are shown. **(B)** Densitometric analysis of Rab10 phosphorylation, as shown in A. The levels of phospho-Rab10 were normalized by those of Rab10. Relative values compared to control (non-target siRNA, CQ (-)) were shown. Data represent mean ± SEM. n = 4, \*\*\*\**P* < 0.0001, ns: not significant, two-way ANOVA with Dunnett’s test. **(C)** Immunoblot analysis of lysates from HEK293 cells stably expressing 3×FLAG-LRRK2 that were additionally transfected with or without V5-Rab29, combined with or without chloroquine exposure for 3 hrs. Representative pictures analyzed for phospho-Thr73-Rab10, Rab10, phospho-S1292-LRRK2, FLAG, V5 and α-tubulin are shown. **(D, E)** Densitometric analysis of Rab10 phosphorylation normalized by total Rab10 levels (D) and Ser1292 autophosphorylation of LRRK2 normalized by total LRRK2 levels (E), as shown in C. Relative values compared to control (mock transfectant) were shown. Data represent mean ± SEM. n = 4, \**P* < 0.05, \*\**P* < 0.01, \*\*\**P* < 0.001, ns: not significant, one-way ANOVA with Tukey’s test. **(F)** A model of two-step activation of LRRK2 phosphorylation of Rab10 by Rab29 and lysosomotropic drugs. Rab29-mediated recruitment of LRRK2 onto organelle membranes enhances LRRK2 enzymatic activity, as monitored by pSer1292 autophosphorylation. Lysosomal overload by drug accumulation facilitates organelle trafficking or fusion, resulting in an increase in the molecular proximity between LRRK2 and substrate Rab GTPases that were otherwise localized on distinct organelle membranes.

We next examined whether the Rab29-mediated mechanism is sufficient to explain the Rab10 phosphorylation by the lysosomotropic agents. HEK293 cells stably expressing human wild-type LRRK2 were transfected with V5-tagged Rab29 and further incubated with or without chloroquine for 3 hrs. Coexpression of Rab29 increased phosphorylation of both LRRK2-Ser1292 and Rab10-Thr73, whereas additional treatment with chloroquine selectively upregulated the Rab10-Thr73 phosphorylation **(Fig. 6C-6E)**. This result suggested that Rab29 acts upstream of the activation of LRRK2 phosphorylation of Rab10 by lysosomotropic agents. As it has been well demonstrated that Rab29 expression facilitates the translocation of LRRK2 from cytosol to membranes (Gomez et al., 2019; Liu et al., 2018; Purlyte et al., 2018), these results collectively support the two-step activation model, in which LRRK2 enzymatic activity is first enhanced by its membrane translocation by Rab29, then its activity to phosphorylate Rab GTPases is further enhanced by the lysosomal stress that facilitates LRRK2-substrate interactions on membranes **(Fig. 6F)**.

## Discussion

In this study, we revealed that lysosomotropic drugs commonly activate LRRK2 phosphorylation of Rab10 as well as the LRRK2-mediated exocytic release of lysosomal content. Unexpectedly, the upregulation of Rab10 phosphorylation was not attributed to the proportional increase of LRRK2 enzymatic activity, but rather reflected an increase in the molecular proximity between LRRK2 and Rab10. This point of view is of particular importance to interpret the results of altered Rab phosphorylation by LRRK2, as the phosphorylation in the switch II region of Rab, especially Rab10, has been regarded as a biomarker to measure the activity of LRRK2 kinase both in experimental settings and clinical studies.

We also revealed that Rab10 phosphorylation was specifically upregulated by pharmacological induction of lysosomal stress, but not other types of cellular stresses. There may be a couple of different mechanistic explanations for this observation. First, endogenous LRRK2 has the potential to accumulate more on overloaded lysosomes than on other organelles (Fig. 2A) (Eguchi et al., 2018). Second, lysosomal overload may cause the aberrant membrane trafficking or organelle fusion more readily than other cellular stresses. LRRK2 substrates Rab10 and Rab8 are normally localized to the endosomal membranes including recycling and late endosomes, in contrast to LRRK2 that resides on the Golgi, and the increased membrane trafficking or fusion is expected to facilitate the assembly of LRRK2 and Rab on the same membranes. We observed that monensin as well as chloroquine induced the upregulation of Rab10 phosphorylation more potently than bafilomycin A1 or NH4Cl, which may be due to the stronger potential of the former two drugs to increase the lysosomal osmotic pressure (Florey et al., 2015), although the pharmacological effect of monensin is not fully understood. The lysosomal overload stress may not only be brought on by artificial conditions such as chloroquine treatment but occur under physiological conditions, as LRRK2-coated enlarged lysosomes were observed in a small but significant population of healthy cells, i.e., without chloroquine treatment (Eguchi et al., 2018).

It should be noted that the concentrations of most of the drugs used in this study are much higher than those used in clinical applications. The only exceptions are those with some local anesthetic drugs, *e*.*g*., lidocaine and procainamide used in this study, because acute dosing of these local anesthetics at a concentration of clinical use has been shown to cause massive cytoplasmic vacuolization, which has been documented in a variety of studies including human clinical studies (biopsy samples of epidermal and dermal cells) (Cazes et al., 2007; Hoss et al., 1999; Vallance et al., 2004), animal experiments (ocular tissues) (Anderson et al., 1999; Atilla et al., 2003) and *in vitro* studies using primary cultured cells (human skin fibroblasts, rabbit smooth muscle cells) (Bawolak et al., 2010; Michalik et al., 2003; Morissette et al., 2004). Also in such studies using primary cells, the vacuole formation by these local anesthetics at this concentration has been shown to be fully reversible (Bawolak et al., 2010; Michalik et al., 2003), suggesting that the concentration is below toxic thresholds. However, it might be misleading to assume that the administration of these drugs in patients immediately causes a type of side effect as a result of aberrant upregulation of Rab10 phosphorylation, since the effect of such upregulation is expected to occur in extremely time- and region-specific manners.

We also confirmed that LRRK2 kinase activity was a key regulator of the exocytic release of lysosomal content under lysosomal overload stress. Although the phenomenon of the stress-induced lysosomal exocytosis has long been documented (Gonzalez-Noriega et al., 1980; Pohlmann et al., 1984; Takano et al., 1984; Tapper and Sundler, 1990), little is known about its molecular mechanism. LRRK2 inhibitor treatment significantly reduced the exocytic cathepsin B release upon lysosomotropic drug exposure, but the reduction was not necessarily complete. It may be that nonphosphorylated Rab10 or other substrate Rabs are also capable of regulating this release process, albeit less efficiently; alternatively, other as yet unidentified factors that are not related to LRRK2 kinase activity may partly be involved in this process. Regarding Rab GTPases that are responsible for the cathepsin release upon lysosomal stress, we previously identified Rab10 as the main mediator downstream of LRRK2 (Eguchi et al., 2018), but the involvement of phosphorylation of other LRRK2 substrate Rabs, *e*.*g*., Rab3 or Rab35, should also be examined as these Rabs have been implicated in the lysosomal exocytosis or exosome release.

It would be tempting to speculate if this release mechanism were involved in the propagation of α-synuclein or other pathogenic proteins in neurodegenerative diseases, especially of PD (Baba et al., 1998; Fujiwara et al., 2002). The accumulation of overloaded lysosomes is an age-dependent histological alteration commonly observed in aged tissues, where waste products such as lipofuscin granules are accumulated (Carmona-Gutierrez et al., 2016; Cuervo and Dice, 2000; Gomez-Sintes et al., 2016). Specifically, it has also been reported that LRRK2 is aberrantly localized to endolysosomal compartment in Lewy body disease brains (Higashi et al., 2009). In view of our hypothesis, one may argue that these represent the accumulation of LRRK2-coated overloaded lysosomes. Recent studies pointed to the close relationship between LRRK2 and another PD risk factor glucocerebrosidase (GCase), a lysosomal enzyme that hydrolyzes glucosylceramide to ceramide and glucose (Sanyal et al., 2020; Yahalom et al., 2019; Ysselstein et al., 2019). Notably, it has been shown that LRRK2 kinase activity negatively regulates GCase activity through Rab10 (Ysselstein et al., 2019). Taken together with our observations, a positive feedback loop mechanism that may lead to PD pathophysiology could be proposed; that is, the reduction in GCase activity causes lysosomal overload that enhances LRRK2 phosphorylation of Rab10, which in turn decreases the GCase activity and further impairs the lysosome integrity. Collectively, it would be possible to assume that lysosomal overload is accelerated in aged human brains at the risk of developing PD, which may potentially be accompanied by the upregulation of LRRK2 phosphorylation of Rab10. Persistence of this cellular alteration may then lead to the aberrant exocytic release of lysosomal content or defective maintenance of lysosomal homeostasis, finally leading to the neuronal death observed in PD. Although this pathogenetic process is one of the plausible mechanisms in sporadic PD, the process in familial PD with LRRK2 mutations might be different; pathogenic mutant LRRK2 is known to have a higher kinase activity intrinsically and therefore may cause the above-described pathogenetic process through hyperphosphorylated Rab10 by bypassing the lysosomal overload. In other words, the lysosomal overload may not occur as a consequence caused by LRRK2 mutations in familial PD associated with LRRK2.

Preventing the lysosomal overload would be regarded as a possible way to block LRRK2-induced neurodegeneration. Thus, it may be crucial to establish a novel therapeutic strategy to maintain lysosomal activity so that neurons in PD and other neurodegenerative disorders could eliminate waste products accumulated in lysosomes that may lead to neuronal degeneration.

## Materials and Methods

### Plasmids and siRNA

The plasmids encoding 3×FLAG-tagged full-length human LRRK2 (wild-type, G2019S, R1441C, Y1699C) were generated previously (Fujimoto et al., 2018). The plasmid encoding V5-Rab29 was generated by transferring rat Rab29 cDNA sequence in pEGFP-C1 vector (MacLeod et al., 2013) to pcDNA3.1/nV5-DEST vector via subcloning into pENTR/D-TOPO followed by LR reaction. siRNA mixtures for mouse LRRK2 and Rab29, which contained 4 different siRNAs per target, were purchased from GE Dharmacon (siGENOME SMARTpool).

### Reagents

The following reagents causing cellular stresses were used at final concentrations as indicated: chloroquine (50-100 μM, Sigma), bafilomycin A1 (100 nM, Wako), ammonium chloride (10 mM, Wako), monensin (100 μM, Cayman), vacuolin-1 (500 nM, Santa Cruz), Torin 1 (1 μM, Cell Signaling), CCCP (50 μM, Sigma), tunicamycin (10 μg/ml, Cayman), rotenone (500 nM, Sigma), lipopolysaccharide (5 μg/ml, Sigma). Clinically used lysosomotropic drugs were as follows: chlorpromazine (Tokyo Kasei Kogyo), thioridazine (Cayman), nortriptyline (Tokyo Kasei Kogyo), fluoxetine (Tokyo Kasei Kogyo), imatinib (Tokyo Kasei Kogyo), gefitinib (Tokyo Kasei Kogyo), tamoxifen (Sigma), lidocaine (Fujifilm), procainamide (Fujifilm), nifedipine (Wako). The final concentrations of these drugs were indicated in Table 1. GSK2578215A was obtained from Sigma.

### Antibodies

The following antibodies were used: anti-LRRK2 [MJFF2 (c41-2)] (Abcam), anti-LRRK2 [N138/6] (NeuroMab, for immunoprecipitation), anti-phospho-Ser1292 human LRRK2 [ab203181] (Abcam), anti-phospho-Ser935 LRRK2 [5099-1] (Epitomics), anti-Rab10 [D36C4] (Cell Signaling, for western blotting), anti-Rab10 [4E2] (Abcam, for proximity ligation assay), anti-phospho-Thr73 Rab10 [MJF-R21 (ab230261)] (Abcam), anti-Rab8a rabbit monoclonal antibody (mAb) [EPR14873 (ab188574)] (Abcam, for western blotting), anti-Rab8 mouse mAb [610845] (BD Transduction Laboratories, for proximity ligation assay), anti-phospho-Thr72 Rab8A [MJF-R20 (ab230260)] (Abcam), anti-mouse LAMP1 [1D4B] (Bio-Rad, for immunocytochemistry), anti-LAMP1 cytosolic tail [L1418] (Sigma, for proximity ligation assay), anti-FLAG [M2] (Sigma), anti-α-tubulin [DM1A] (Sigma), anti-cathepsin B [D1C7Y] (Cell Signaling), anti-LBPA [6C4] (Echelon), anti-EEA1 [#2411] (Cell Signaling). The rabbit polyclonal antibody against Rab29 was generated previously (Aizawa and Fukuda, 2015).

### Cell culture, transfection and drug treatment

HEK (human embryonic kidney) 293 cells, SH-SY5Y cells and 3T3-Swiss albino cells were cultured in Dulbecco’s modified Eagle’s medium (DMEM) supplemented with 10% (v/v) fetal bovine serum and 100 units/mL penicillin, 100 μg/mL streptomycin at 37°C in 5% CO2 atmosphere. RAW264.7 cells (obtained from ATCC) were cultured on culture dishes for suspended cells (Sumitomo Bakelite Co.) under the same conditions with HEK293 cells. RAW264.7 cells were always pretreated with IFN-γ (15 ng/ml, Cell Signaling) for 48 hrs before each assay. Bone marrow-derived primary macrophages were extracted from mouse femurs and tibias and then cultured as described previously (Eguchi et al., 2018). The experiment with mice was performed in accordance with the regulations and guidelines of the University of Tokyo and approved by the institutional review committee. HEK293 cells stably expressing human LRRK2 (WT) were generated by transfection of 3×FLAG-tagged LRRK2 (WT) plasmid followed by pharmacological selection using an antibiotic G-418 (Roche). Transfection of plasmids and siRNA was performed using Lipofectamine 3000 (Thermo Fisher Scientific) and Lipofectamine RNAiMAX (Thermo Fisher Scientific), respectively, according to the manufacturer’s protocols. For transfection of LRRK2, plasmids were diluted five-fold with mock vector (p3×FLAG-CMV-10) upon transfection to reduce the extent of overexpression. For RNAi experiments, cells were analyzed 72 hrs after siRNA transfection. Lysosomotropic drugs at the concentrations as listed in Table 1 were added to cells by medium replacement, and PBS was added for controls.

### Immunoblot analysis

Cells were washed with PBS on ice and lysed in a lysis buffer containing 50 mM Tris HCl pH 7.6, 150 mM NaCl, 1% (v/v) Triton X-100, Complete EDTA-free protease inhibitor cocktail (Roche), and PhosSTOP phosphatase inhibitor cocktail (Roche). Lysates were centrifuged at 17,800 × *g* for 5 min at 4°C and supernatants were mixed with NuPAGE LDS Sample Buffer (4×) buffer (Thermo Fisher Scientific) containing 1% (v/v) 2-mercaptoethanol. For SDS-PAGE, samples were loaded onto Tris-glycine gels, electrophoresed and transferred to PVDF membranes. Transferred membranes were blocked and incubated with primary antibodies and then with HRP-conjugated secondly antibodies (Jackson ImmunoResearch). Protein bands were detected by LAS-4000 (Fujifilm). The integrated densities of protein bands were calculated using ImageJ software (NIH).

### Measurement of cathepsin B and LDH in media

Media from RAW264.7 cells were collected and centrifuged at 300 × *g* for 5 min, and the supernatants were subjected to immunoblotting and LDH assay. For immunoblotting, the supernatants were mixed with NuPAGE LDS Sample Buffer (4×) buffer (Thermo Fisher Scientific) containing 1% (v/v) 2-mercaptoethanol before loading onto Tris-glycine gels. The activity of LDH in the supernatants were measured using Cytotoxicity Detection Kit (Roche Applied Science) according to the manufacturer’s protocol.

### Immunoprecipitation

RAW264.7 cells pretreated with IFN-γ treatment for 48 hrs were treated with CQ (100 μM, 3 hrs) and then lysed in ice-cold lysis buffer (50 mM Tris-HCl pH 7.6, 150 mM NaCl, 0.5%(v/v) Triton X-100, protease inhibitor cocktail (cOmplete EDTA-free, Roche) and PhosSTOP (Roche)) followed by centrifugation at 20,800 × *g* for 5 min at 4 °C. 500 µL of the cleared lysates were added with 20 µL Recombinant Protein G Agarose (Invitrogen) and rotated for 30 min at 4 °C to preclear. Lysates were then added with 10 µL Recombinant Protein G Agarose and 2 µL antibodies, and rotated for 3 hrs at 4 °C. Immunocomplexes were then precipitated by centrifugation and washed three times with 700 µL of the PBS. Bound proteins were eluted by boiling the beads in 40 µL of NuPAGE LDS Sample Buffer (Thermo Fisher Scientific) for 15 min. For immunoblotting, 15 µL of the IP samples were separated by Tris-glycine SDS-PAGE gels. As for “input” samples, 15 µL of cell lysates were loaded alongside the IP samples. Anti-LRRK2 antibody [N138/6] and anti-mouse IgG antibody were used for immunoprecipitation.

### Immunocytochemistry

Immunofluorescence analysis was performed based on our previous methods (Eguchi et al., 2018; Kuwahara et al., 2016). Cells cultured on coverslips were fixed with 4% (w/v) paraformaldehyde for 20 min, followed by immersion in 100% EtOH at -20°C. 100% EtOH treatment was necessary for the staining with an anti-LRRK2 antibody clone c41-2 (MJFF2). Samples were washed with PBS and then permeabilized and blocked with 3% (w/v) BSA in PBS containing 0.5% Triton X-100. Primary antibodies and corresponding secondary antibodies conjugated with Alexa Fluor dyes (Thermo Fisher Scientific) were diluted in the blocking buffer, and samples were incubated with antibody solutions. The samples were imaged using a confocal microscope (SP5, Leica). Image contrast and brightness were adjusted using Photoshop 2020 software (Adobe). For the quantitative analysis of LRRK2-LAMP1 colocalization, Manders’ coefficient on the fluorescence images was calculated using the ImageJ JACoP plugin, with the auto-threshold method.

### Proximity ligation assay (PLA)

Proximity ligation was performed using Duolink PLA probes (Sigma) and In situ Detection Reagents Orange (Sigma) according to the manufacturer’s protocols. The essential procedures are as follows. Cells were PFA-fixed, permeabilized and blocked as described for immunocytochemistry. Cells were then incubated with the combination of two primary antibodies of interest diluted in 3% BSA/PBS for 3 hrs. For labeling of LAMP1, rabbit anti-LAMP1 antibody that recognizes its cytosolic tail (L1418, Sigma) was used. After washing with PBS, cells were incubated with anti-mouse PLUS and anti-rabbit MINUS probes for 1 hr at 37°C. Cells were washed with Buffer A (0.01M Tris, 0.15M NaCl, 0.05% Tween 20, pH7.4) and reacted with the ligase in 1× ligation buffer for 30 min at 37°C. After washing with Buffer A, cells were incubated with the polymerase in 1× amplification buffer for 100 min at 37°C in a pre-heated humidity chamber. Cells were then washed with Buffer B (0.2M Tris, 0.1M NaCl, pH7.5) and incubated with DRAQ5 (Biostatus) for 30 min to stain nuclei. Cells were washed again with Buffer B and 0.01× Buffer B followed by encapsulation. The samples were imaged using a confocal microscope (SP5, Leica). For quantitative analysis, pictures containing 100-200 cells were taken by 40× objective lens of confocal microscope, and five representative fields from two independent experiments were collected. PLA particles as well as nuclei in each picture were automatically counted by ImageJ software (NIH) using the Analyze Particles mode, and PLA particles in each cell were calculated by dividing total number of PLA particles by that of nuclei.

### Statistics

The statistical significance of difference in mean values was calculated either by unpaired, two-tailed Student *t* test, one-way ANOVA or two-way ANOVA, as indicated. *P* values less than 0.05 were considered statistically significant. No exclusion criteria were applied to exclude samples or animals from analysis.

## Acknowledgements

The authors thank all members in Iwatsubo’s lab, especially Drs. Tadafumi Hashimoto, Tomoko Wakabayashi and Kaoru Yamada for helpful discussions. This work was supported by JSPS KAKENHI Grant Numbers 16K07039, 19K07816, 20H00525 and 20J12819.

## Author contributions

T. Ku. designed the research, performed cell culture experiments and wrote the manuscript. K. F., T. Ko., M. S. and G. Y. performed cell culture experiments. T. E. and M. F. provided the idea for the research and prepared materials. T. I. provided intellectual input for the research and wrote the manuscript with T. Ku.

## Abbreviations

LRRK2: Leucine-rich repeat kinase 2
PD: Parkinson’s disease
CQ: chloroquine
CCCP: carbonyl cyanide m-chlorophenylhydrazone
LPS: lipopolysaccharide
WT: wild-type
GSK: GSK2578215A
LDH: lactate dehydrogenase
LAMP1: lysosomal-associated membrane protein 1
Cat B: cathepsin B
LBPA: lysobisphosphatidic acid
EEA1: early endosomal antigen 1
PLA: proximity ligation assay
GCase: glucocerebrosidase
EtOH: ethanol
BSA: bovine serum albumin
PBS: phosphate-buffered saline

