## supplementary figures for "Roles of lysosomotropic agents on LRRK2 activation and Rab10 phosphorylation"

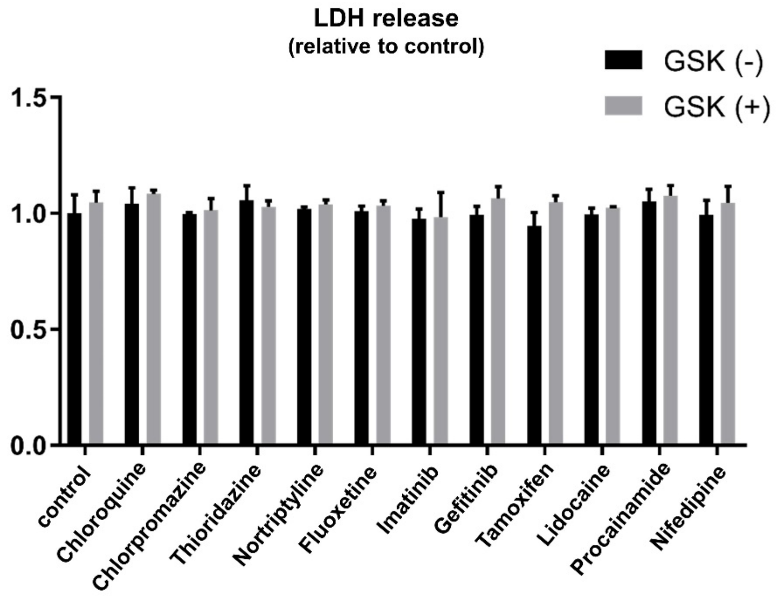


**Supplementary Fig. 1. Treatment with lysosomotropic drugs does not cause cell death.**

LDH activity in media from RAW264.7 cells treated with the indicated drugs with or without LRRK2 kinase inhibitor GSK2578215A (GSK). The activities relative to control-GSK (-) are shown. Data represent mean ± SD of duplicate measurements.

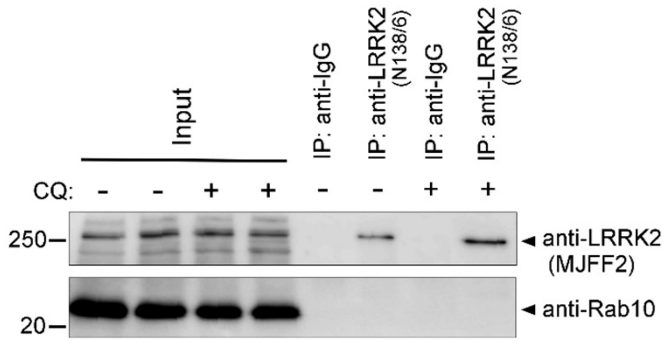


**Supplementary Fig. 2. Lack of stable interaction between LRRK2 and Rab10 in cells treated with chloroquine.**

Endogenous LRRK2 was immunoprecipitated from RAW264.7 cells using anti-LRRK2 monoclonal antibody N138/6. Anti-mouse IgG antibody was used as a negative control. Endogenous Rab10 was not co-precipitated with LRRK2, even after treatment with chloroquine.
